# Electron-counting in MicroED

**DOI:** 10.1101/2023.06.29.547123

**Authors:** Johan Hattne, Max T. B. Clabbers, Michael W. Martynowycz, Tamir Gonen

**Affiliations:** Howard Hughes Medical Institute, University of California, Los Angeles, CA 90095; Department of Biological Chemistry, University of California, Los Angeles, CA 90095; Department of Physiology, University of California, Los Angeles, CA 90095

**Keywords:** Electron-counting, MicroED, Microcrystal electron diffraction, Cryo-EM

## Abstract

The combination of high sensitivity and rapid readout makes it possible for electron-counting detectors to record cryogenic electron microscopy data faster and more accurately without increasing the exposure. This is especially useful for MicroED of macromolecular crystals where the strength of the diffracted signal at high resolution is comparable to the surrounding background. The ability to decrease the exposure also alleviates concerns about radiation damage which limits the information that can be recovered from a diffraction measurement. However, the dynamic range of electron-counting detectors requires careful data collection to avoid errors from coincidence loss. Nevertheless, these detectors are increasingly deployed in cryo-EM facilities, and several have been successfully used for MicroED. Provided coincidence loss can be minimized, electron-counting detectors bring high potential rewards.

## Introduction

Microcrystal electron diffraction (MicroED) is a technique in cryogenic electron microscopy (cryo-EM) that has proven effective for determining structures from crystalline samples that are only a few hundred nanometers thick (Shi *et al*., 2013). The method has significant advantages in structural biology because growing large and well-ordered crystals is often the bottleneck in crystallographic structure determination. This advantage stems from the larger cross-section and more favorable ratio of elastic to inelastic interactions of electrons with matter (Henderson, 1995), which means that even for minuscule crystals a low electron flux can provide the necessary information to determine the structure at atomic resolution (Martynowycz *et al*., 2022).

The resolution of the measurable data from a diffracting crystal is limited by the relative intensity in the reflections compared to the surrounding background. The intensity is proportional to the number of scattered electrons, which increases with the number of unit cells illuminated by the beam, so the diffracted intensity from a small crystal with fewer unit cells is correspondingly weaker. A faint reflection may not be discernible from the contribution of the solvent in or around the crystal, lattice disorder, or the Poisson noise of the scattering process. High-resolution features are particularly susceptible to this limitation because the spots from which they derive are inherently weaker than the low-resolution reflections near the center of the diffraction pattern. Therefore, sensitive electron detection devices are needed to accurately record the pixel values of the diffraction pattern to the highest resolution possible.

In the past, weak reflections in MicroED have typically been recovered by increasing the flux. This also leads to higher energy absorption by the sample, resulting in more radiation damage (Hattne *et al*., 2018). This dose-damage trade-off is not unique to MicroED, but is a concern for all modalities of structural biology, particularly those dealing with very small samples (Holton and Frankel, 2010; Yamamoto *et al*., 2017). Increasing the flux boosts the signal at high resolution but also amplifies the intensity of the strong spots at low resolution. This is not an issue for integrating detectors with a high dynamic range but can be problematic for electron-counting detectors, which are limited in their ability to distinguish individual electrons at high flux.

Electron-counting cameras can measure very low electron counts at high quantum efficiency. Unlike scintillator-based CCD or CMOS detectors that integrate the accumulated secondary effects, electron-counting devices rely on direct detection. The charge separation from primary electrons incident in the sensor are compared to set thresholds and individually registered, allowing single-electron events to be represented. This means that the flux can be reduced without compromising the accuracy of the measurement. However, counting cameras face a significant challenge when dealing with strong reflections (Leonarski *et al*., 2018). After an electron hits a pixel, the read-out electronics must reset before the next electron can be counted, lest two electrons striking the same pixel in rapid succession will be recorded as a single event. Such coincidence loss causes the detector response to become nonlinear and disrupts data reduction, which is based on the premise that the integrated intensity is proportional to the number of scattered electrons.

The limited count-rate of direct electron detectors poses a challenge for macromolecular MicroED. But with careful data collection and analysis, these detectors can provide data to higher resolution at a lower total exposure. For weakly diffracting crystals of radiation-sensitive molecules, they may prove critical for the success of structure solution.

## Results and discussion

### Coincidence loss

Direct electron detectors operate at high internal frame rates (250 Hz for the Falcon 4, 320 Hz for the Falcon 4i, 400 Hz for the K2 Summit, and 1500 Hz for the K3). In counting mode, at most one electron can be registered on a given pixel per internal frame and subsequent electrons on the pixel during the readout cycle are ignored. Thus, the pixel values in an internal frame are inherently binary. To increase the information content per data transfer, the camera system sums batches of consecutive frames. Summation reduces the effective frame rate (∼35 Hz for the Falcon 4, 40 Hz for the K2 Summit, and >75 Hz for the K3) but does not degrade the accuracy of the pixel values since both Landau and readout noise are negligible (Battaglia *et al*., 2009). Because summation also increases the effective dynamic range for each frame, individual electrons can still be discerned in the recorded data (*Figure 1*).

**Figure 1:**
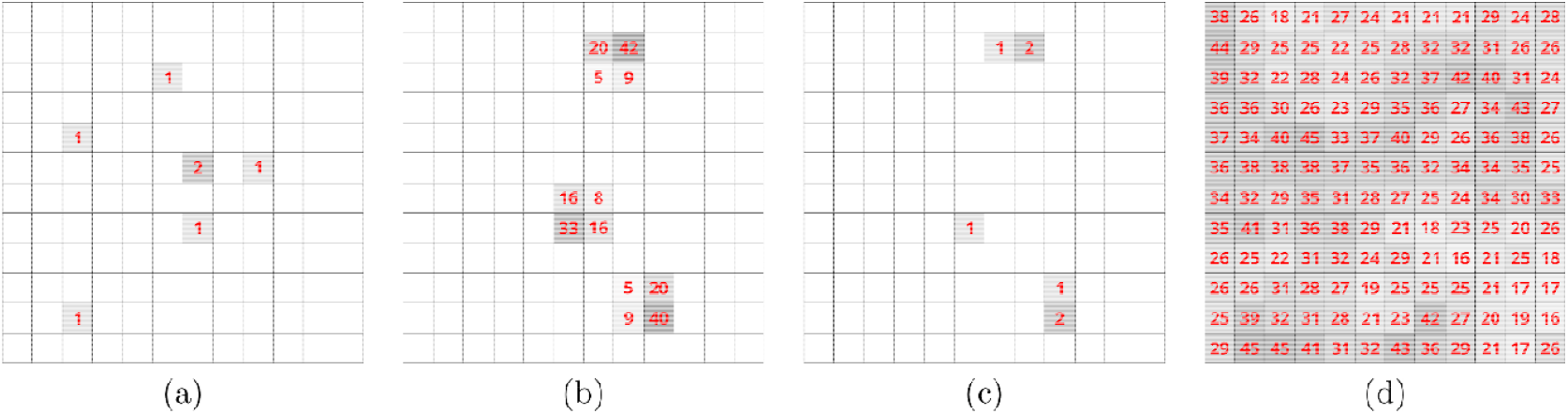
Single-electron events are typically recorded on a single pixel on a direct electron-detector. (a) Sum of 20 internal frames over 13⍰μs on the K3. (b) MRC output of 35 summed internal frames over 140⍰μs on the Falcon 4. (c) The pixels in (b) after sharpening, which reduces the artificial spread introduced by the MRC output from the Falcon 4. (d) For comparison, output from the Falcon 3 in integrating mode. All panels show a 12×12 pixel area at ∼2⍰Å resolution in diffraction patterns from proteinase K. The unlabeled pixels in (a), (b), and (c) are zero.

Coincidence loss happens before the dynamic range of a pixel has been exhausted. Assuming the arrival time of *N* electrons on *M* internal frames is randomly distributed, such that each frame in a batch is equally likely to see an electron, the probability of zero coincidence loss, *p*_*M*_(*N*), may be recursively modelled as

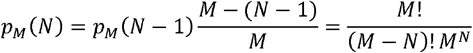

where *p*_*M*_(1) = 1. This is a simplification, because the actual arrival time is not randomly distributed, but for sufficiently small *M*, the error may be acceptable. For example, for *M* = 10 internal frames, the probability of observing *N* = 8 electrons without coincidence loss is exceedingly small, *p*_10_(8) = 0.018 (*Figure 2*). For moderate coincidence loss, postprocessing in either hardware (Trueb *et al*., 2015) or software (Gallagher-Jones *et al*., 2019) can model some of the information lost due to electron pile-up, but the gain in accuracy may not warrant the added complexity as pixels must be tracked over several frames.

**Figure 2:**
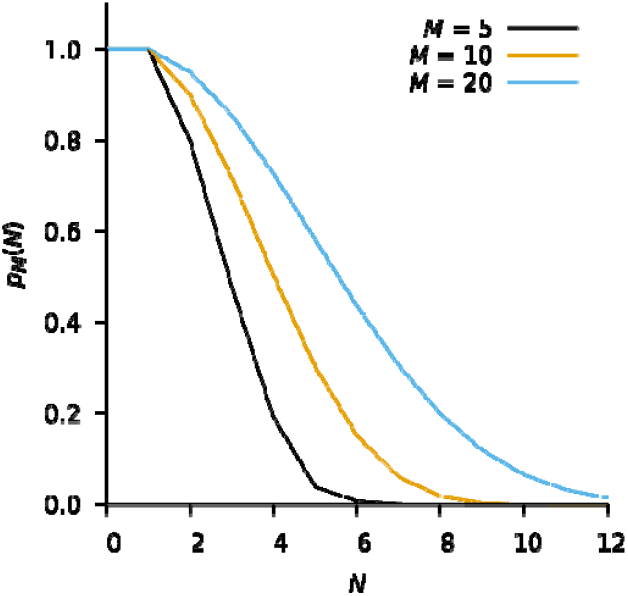
The modelled probability of zero coincidence loss on 5 (black), 10 (orange), and 20 (blue) internal frames. Increasing the exposure time, or equivalently, increasing the number of internal frames, M, increases the probability of zero coincidence loss for a given number of incident electrons, but also implies more incident electrons.

The only way to avoid detrimental corruption of the pixel values while collecting an electron-counted dataset on a given camera is to reduce the exposure. To avoid sacrificing the weak high-resolution reflections, data can be collected in two passes (Dauter, 2017): one pass aimed at recording the strongest low-resolution reflections and another pass for the high-resolution reflections. The exposure during the low-resolution pass is minimized to avoid coincidence loss and radiation damage. The exposure is significantly higher for the high-resolution pass to capture the weak reflections without concern for pile-up in the low-resolution data.

In practice, some coincidence loss is inevitable. For overexposed samples, all pixel values integrated for a reflection are ultimately capped at the maximum count determined by the number of internal frames summed in the camera. Measures of internal consistency such as *CC*_1/2_ and *R*_merge_ may not detect degradation of data quality due to coincidence loss because the integrated intensities tend to become more similar as intensities are only underestimated, but never overestimated. Instead, detrimental pile-up may be diagnosed from poor correlations between observed and calculated low-resolution reflections during structure refinement (Clabbers *et al*., 2022).

### Fine-slicing and bandwidth concerns

The high sensitivity and high frame rate allow counting data to be collected in much finer slices than previously possible with scintillator-based cameras. Fine slicing of reciprocal space produces a large amount of data in a short time but captures the scattering process at much greater detail and allows for more accurate modelling of spot profiles (Pflugrath, 1999). This information can be useful in refining the crystal structure and understanding its properties.

The rotation range of the finest measurable slice is determined by the effective frame rate of the camera and the slowest permissible rotation speed of the sample stage. In practice, the smallest rotation range is bound by the camera system’s memory buffers and network bandwidth because data must be transferred off the camera system in near-real time. However, even after internal summation, most pixel values in an electron-counted MicroED dataset are zero (*Figure 3*). Standard compression algorithms tend to perform well on such sparse data and the *MicroED tools* (see section Availability) can now decompress and convert such files on the fly. Alternatively, emerging event-based image formats make more efficient use of computational resources in the camera system (Gallego *et al*., 2022) and allow internal summation to be skipped altogether. While event-based formats are well-suited to store and transmit electron-counted data, they are not yet widely supported by pipelines for diffraction data reduction.

**Figure 3:**
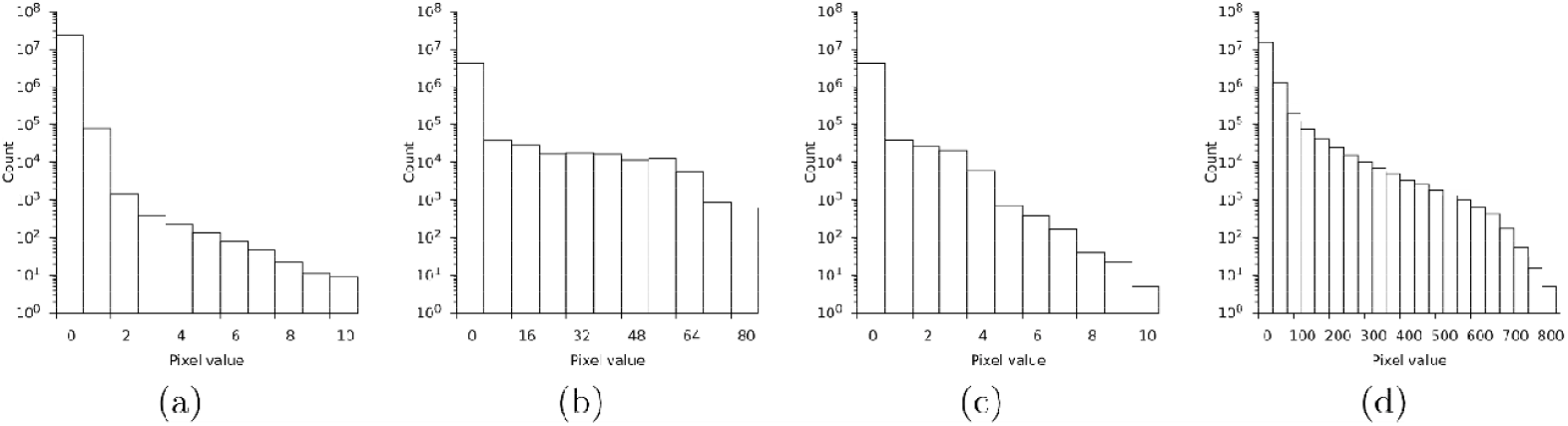
Logarithmic histogram of pixel values on a single output frame. (a) Sum of 20 internal frames on the K3. (b) MRC output of 35 summed internal frames on the Falcon 4. (c) The Falcon 4 batch after sharpening. (d) The Falcon 3 in integrating mode for comparison.

### Image analysis

Electron-counting data have a noticeably different appearance from data recorded on an integrating, indirect detector (*Figure 1*). Because neither a scintillator nor fiberoptic coupling are needed the point-spread is essentially zero and the size of a reflection is almost exclusively determined by the crystal size, mosaicity, and beam divergence. The distinctly discrete nature of electron-counting diffraction images demands different analysis methods. This is not unique to MicroED data. For instance, *Adxv* has options for enhancing spots on very weak backgrounds, and integration in *XDS* (Kabsch, 2012) is often improved by using settings originally devised for photon-counting X-ray detectors (Clabbers *et al*., 2022).

The cameras used in cryoEM are generally developed for single-particle analysis, where the goal is to obtain the highest possible spatial resolution of images of single macromolecules. Recent models can be operated in superresolution mode, which increases spatial resolution by calculating the point of impact of an electron on the detector surface to subpixel precision. Superresolution is not as important for MicroED, where images are typically binned further to reduce processing time and provide sharper diffraction patterns. With the Falcon 4, some superresolution information is retained in the MRC output by convoluting the raw data with different kernels depending on the sub-pixel position of the electron event. This convolution blurs the electron event over a small area, which makes the distribution of pixel values more uniform and complicates recovery of single-electron events during electron diffraction data reduction (*Figure 3*). At high resolution, where electron events are well separated, single electron events can be approximated by deconvoluting with matching kernels (*Figure 1*). Sharpening at low resolution is difficult, because the electron events are closely spaced, and the tails introduced by the convolution overlap. Nevertheless, sharpening the Falcon 4 data can partially recover the overall pixel statistics of a raw electron-counted diffraction pattern.

To ensure uniform response across the detector surface, electron-counting data need to be gain-corrected. The most recent *MicroED tools* (see section Availability) can apply a gain correction while summing and converting the data from the camera system. Using data directly from the camera system is important, as any intermediate conversion steps carry the risk of introducing unwanted transformations. Since unit gain images, where each electron increases the corresponding pixel value by one, are impractical for integer-based image formats such as SMV, pixel values can additionally be scaled to avoid excessive loss of precision. For example, multiplication by 32 will preserve five bits of the fractional precision, which is usually sufficient.

## Conclusion

Fast-counting detectors have been used in X-ray crystallography for many years (Broennimann *et al*., 2006), where they have enabled shutterless collection of finely sliced rotation diffraction data. Their more recent use in cryo-EM is largely motived by their high frame rate, which enables image processing algorithms to correct for beam-induced motion which otherwise introduces blurring as the sample moves during the exposure (Li *et al*., 2013). In contrast, reciprocal space as captured on a diffraction pattern in MicroED is relatively invariant to translation. A fast-framing detector with negligible dead time allows data to be collected in continuous rotation mode without introducing gaps between frames (Nannenga *et al*., 2014). Continuous rotation under constant exposure avoids the need for synchronization between stage ramp-up, beam shuttering, and readout and has become common in other modes diffraction data collection as well (Powell, 2017).

The use of electron counting has greatly improved the accuracy of MicroED data. In early work with a scintillator-based camera, practical considerations forced sacrifices in accuracy. To enable continuous rotation data collection the camera was configured in rolling shutter mode (Stumpf *et al*., 2010). In this mode, the TemCam-F416 could record data in a single uninterrupted sweep, but had no time for *e*.*g*. correlated double-sampling, and intensity modelling was required to recover high-resolution information lost to truncation (Hattne *et al*., 2016). The processing pipeline with a counting direct electron detector is both simpler and provides more accurate data. A striking illustration is the successful phasing of two different macromolecules without reference to their known structures (Martynowycz *et al*., 2022). Such *ab initio* phasing is critically dependent on the data quality. Another example is provided by the structure of a protoglobin variant from crystals that had long resisted solution by synchrotron X-ray crystallography (Danelius *et al*., 2023).

Due to their higher sensitivity, direct electron detectors allow for lower flux and shorter exposure times without compromising the quality of the data. This reduces the total dose absorbed by the sample, which in turn limits radiation damage (Hattne *et al*., 2018, 2019). The ability to collect useful data using fewer electrons has been a hallmark of MicroED since its inception, and the total exposure during data collection steadily decreased as the method has evolved (*Table 1*).

**Table 1:** Merging and refinement statistics from proteinase K over time. Both resolution and I/σ(I) have generally improved despite the decreased total exposure. These advances are directly affected by developments in detector technology.

| Camera | TemCam-F416 (6cl7) (Hattne *et al.*, 2018) | Falcon 3 (6pu4) (Hattne *et al.*, 2019) | CetaD (7jsy) (Martynowycz and Gonen, 2021) | Falcon 4 (7skx) (Martynowycz *et al.*, 2022) | Falcon 4 (8fyq) (Martynowycz *et al.*, 2023) |
| --- | --- | --- | --- | --- | --- |
| Resolution (Å) | 20.74–1.71 | 27.64–2.10 | 43.32–1.78 | 43.35–1.50 | 19.77–1.40 |
| $I / \sigma(I)$ | 3.8 | 3.1 | 7.8 | 5.7 | 7.6 |
| $CC_{1/2}$ | 0.95 | 0.91 | 0.97 | 0.99 | 0.99 |
| $R_{\text{work}} / R_{\text{free}}$ | 22.13 / 25.34 | 22.06 / 26.70 | 16.94 / 20.94 | 14.95 / 20.46 | 13.74 / 17.50 |
| Total exposure (e<sup>-</sup> Å<sup>-2</sup>) | 1.8 | 1.3 | 2.4 | 1.0 | 1.0 |

Coincidence loss remains a concern for electron-counted MicroED data. Future detectors may mitigate this problem by further increasing the frame rate and improving electron-detection algorithms. Alternatively, hybrid detectors are being designed to overcome the problem by automatically switching to low-gain integrating mode when the flux exceeds a set threshold (Leonarski *et al*., 2018). As these technologies become available on larger chip sizes, they have the potential to further push the boundaries of MicroED.

## Acknowledgments

We would like to acknowledge Alexis Rohou (Genentech) for fruitful discussions. This study was supported by the National Institutes of Health P41GM136508. Portions of this research or manuscript completion were developed with funding from the Department of Defense grants MCDC-2202-002 and HDTRA1-21-1-0004. Effort sponsored by the U.S. Government under Other Transaction number W15QKN-16-9-1002 between the MCDC, and the Government. The US Government is authorized to reproduce and distribute reprints for Governmental purposes, notwithstanding any copyright notation thereon. The views and conclusions contained herein are those of the authors and should not be interpreted as necessarily representing the official policies or endorsements, either expressed or implied, of the U.S. Government. The PAH shall flowdown these requirements to its subawardees, at all tiers. The Gonen laboratory is supported by funds from the Howard Hughes Medical Institute.

## Availability

The source code for the MicroED tools with multithreaded summation, compression, and gain-correction support is available from https://cryoem.ucla.edu/downloads. The next upcoming snapshot can convert all versions of Gatan’s Digital Micrograph format to either SMV or TIFF.

## Notes

### Competing Interest Statement

The authors have declared no competing interest.

